# Analyzing and Visualizing Distance Measures for Clonal Trees

**DOI:** 10.1101/2025.07.18.665562

**Authors:** Thea Traw, Quoc Nguyen, Layla Oesper, Eric Alexander

## Abstract

Cancer is an evolutionary process, in that a tumor results from a series of genomic mutations that are acquired over time. The history of how a tumor evolved is often represented using a clonal tree: a particular type of rooted tree where vertices represent tumor cell clones and edges represent ancestral relationships between them. In recent years, numerous algorithms have been developed that infer clonal trees from sequencing data. Correspondingly, researchers have also developed distance measures to compare clonal trees. While such distance measures have proven valuable, they are limited in a variety of ways. First off, they usually only utilize one or a few types of specific information when being computed - making it challenging to assess how well they capture other common types of differences between clonal trees. Secondly, their final output is just a single numerical value - meaning that information on what parts of the trees contributed to the distance is not being readily conveyed. In this work, we address both of these issues. We define a set of four classes of tree differences that describe common ways that clonal trees might differ from each other and analyze these classes using three different clonal tree distance measures. We then introduce a visualization tool called VECTr (**V**isual **E**ncodings for **C**lonal **Tr**ees) that provides an enhanced pairwise comparison of clonal trees using the same three distance measures. In particular, VECTr contains three types of visualizations that allow a user to easily identify what parts of a tree contribute to its distance, including information relevant to when the trees differ in a manner that corresponds to one of our four classes of tree differences. We demonstrate the utility of our visualization tool using a variety of different use cases, including application to both simulated and real clonal trees. Code associated with the project is freely available at: https://github.com/quocodile/visualize-clonal-trees

**CCS Concepts:** **Applied computing Molecular evolution; Computational genomics; Computational biology**.

## 1 Introduction

The clonal theory of cancer describes how a tumor is the result of an evolutionary process [20]. Specifically, this theory describes how different sets of somatic genomic mutations (those that occur during a person’s lifetime) are acquired over time in a collection of cells that form a tumor. Every time a cell divides, mutations may appear in the resulting daughter cells. This means that a tumor– which is inherently a heterogeneous collection of cells, each with a collection of mutations–may be described by a particular type of rooted tree, often referred to as a clonal tree. Each vertex in the tree represents a different clone or population of cells that contain the same mutations. Each edge in the tree represents an ancestral relationship between those clones. A better understanding of how a particular tumor evolved has the potential to lead to more targeted therapies and improved chances to avoid relapse [1, 9, 25].

Recent advances in DNA sequencing technologies have led to a revolution in the ways that tumor evolution can be measured and studied [10]. In particular, there has been incredible interest in developing algorithms to infer a patient’s clonal tree from various types of sequencing data [22], with hundreds of algorithms populating the literature (such as [7, 8, 13, 17, 27] and many others). This volume and diversity of methods stems from the many types of data (e.g., bulk and single-cell DNA sequencing) and signals in that data (e.g., single nucleotide variants, copy number aberrations, etc.) that are used for inferring clonal trees, and speaks to the challenge of this endeavor. While the size of these inferred trees is relatively modest (a collection of 43 such trees across 6 different kinds of cancer assembled in [19] were all 30 nodes or fewer), different methods still produce different tree structures, sometimes dramatically so.

One response to the challenge of accurately inferring clonal trees has been the development of specialized distance measures that allow for the comparison of clonal trees (e.g., [6, 12, 14, 15]). These distance measures are used both to compare clonal trees inferred from the same data using different approaches, but also for benchmarking new inference methods on simulated data (in which case a “ground truth” tree is known). However, one challenge is that these distance measures generally utilize only a specific type of information when comparing two clonal trees. For example, parent-child distance [12] only looks at direct relationships between mutations, but ignores all longer range relationships. A broader analysis of how distance measures behave across a variety of commonly occurring or expected differences between clonal trees (not just the ones they were designed to measure) would be beneficial to better understanding the strengths and limitations of these methods.

Another challenge to using existing distance measures for comparing clonal trees is that while most distances do use information across the entirety of both trees to compute the distance, that information is not precisely conveyed in the provided output. Instead, the distance measure simply reports a single numerical value that encompasses all observed differences. While such a reduction can be useful for tasks like benchmarking, where it is important to know which method produced a tree more similar to the ground truth, it lacks important nuance. In addition to knowing which trees are more similar, it is also important to be able to identify precisely which features of those trees contribute to the found distances. This kind of analysis is essential for both improving clonal tree inference methods but also for users of these methods to understand their limitations or biases.

Visualization is a clear choice for encoding information about clonal trees. The most common way to visualize clonal trees has largely been using the standard node-link diagram [8, 13, 17] with associated node labels. While some tree inference methods have included a visualization component as an aside to their methods that infer clonal trees [2, 5], they don’t extensively focus on this component. In recent years, some tools have been developed that allow for different visualizations of clonal trees, many focusing on an alternative to the standard node-link diagram: the fishplot. A fishplot shows information on the inferred prevalence of each clone over time, in addition to the standard parent-child relationships between those clones. Various packages have been created that allow for different variations of fishplots to be created for clonal trees [18, 21, 23]. However, the creation of a fishplot generally requires additional information about the prevalence of different clones at separate timepoints–data that is not always available. Furthermore, fishplots can make it challenging to label mutations associated with different clones and often lack these labels entirely. Furthermore, to our knowledge there is no current work that incorporates information from distance measures along with visualization of clonal trees.

In this paper we define a set of four classes of tree differences that describe common ways that clonal trees might differ from each other and analyze these classes using three different clonal tree distance measures. We then introduce a visualization tool called VECTr (**V**isual **E**ncodings for **C**lonal **Tr**ees) that provides an enhanced pairwise comparison of clonal trees using the same three distance measures. In particular, VECTr contains three types of visualizations that allow a user to easily identify what parts of a tree contribute to its distance, including information relevant to when the trees differ in a manner that corresponds to one of our four classes of tree differences. We demonstrate the utility of our visualization tool using a variety of different use cases, including application to both simulated and real clonal trees.

## 2 Definitions and Background

In this section we provide several key definitions that will be used throughout this paper. First, in Section 2.1 we define a clonal tree, and then in Section 2.2 we describe several distance measures defined for clonal trees.

### 2.1 Clonal trees

A *clonal tree* is a rooted, node-labeled tree that describes the evolutionary history of a tumor. Each vertex in the tree represents a different *clone* or population of cells that contain the same somatic mutations. Each edge in the tree represents an ancestral relationship between those clones. The node labels on the tree indicate the somatic mutations that first arose in the corresponding clone. Unless otherwise indicated, mutations are inherited by all descendant clones. Figure 1 shows both the underlying mutation inheritance pattern (left), as well as their typical presentation as a clonal tree where a mutation is only labeled on the clone where it first appeared (right). The mutation labels on such a clonal tree often indicate the name of the underlying gene that was mutated, rather than a specific genomic position or type of mutation. Clonal trees are distinct from other trees common in the biology literature–namely phylogenetic trees–in several ways: (1) they have labels on all the nodes, rather than just the leaves; (2) multiple labels may appear on any node; and (3) labels are assumed to be inherited by all descendant nodes.

**Figure 1:**
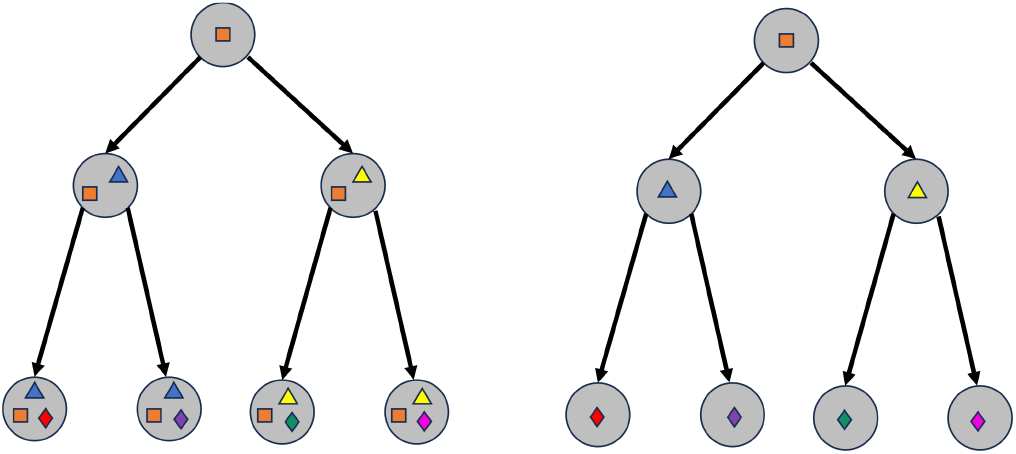
(Left) Diagram showing how mutations (indicated by colored shapes) arise and are inherited by different clones. (Right) The corresponding clonal tree where, for simplicity, the mutations are labeled only at the node in which they were acquired.

### 2.2 Distance measures for clonal trees

In recent years there have been several distance measures developed to compare clonal trees. Specifically, these measures take two clonal trees as input and report a single numerical value indicating how different the two trees are (with a value of 0 typically indicating they are identical). Below we describe three commonly used distance measures on clonal trees: (1) Parent-Child (PC) distance, (2) Ancestor-Descendant (AD) distance, and (3) Distinct-Lineage (DL) distance. For each distance measure, we also specifically describe how it handles multiple mutations first occurring on a single node (a feature of many clonal trees). This important case is not always explicitly described when these distance measures are defined.

#### 2.2.1 Parent-Child distance

Parent-Child (PC) distance counts the number of parent-child mutation pairs that exist in one tree but not the other [12]. I.e., if *PC* (*T)* denotes the set of ordered parent-child pairs observed in tree *T*, then the PC distance between trees *T*_1_ and *T*_2_ is *PC* (*T*_1_, *T*_2_) = |*PC* (*T*_1_)△*PC* (*T*_2_)|, or the size of the symmetric difference () between the associated sets. In Figure 2 the left tree has the following set of ordered parentchild pairs: {(*A, B*), (*B, C*), (*C, D*)} whereas the middle tree has: {(*B, A*), (*B, C*), (*B, D*)}. The symmetric difference between them is {(*A, B*), (*C, D*),(*B, A*),(*B, D*,)} giving a PC distance of 4.

When multiple mutations appear on a single node, each mutation contributes individually to the associated parent-child relationships. For example, if mutations *B* and *C* appear together on a node that is a child of mutation *A*, they would contribute the following parentchild relationships: (*A, B*), (*A, C*).

**Figure 2:**
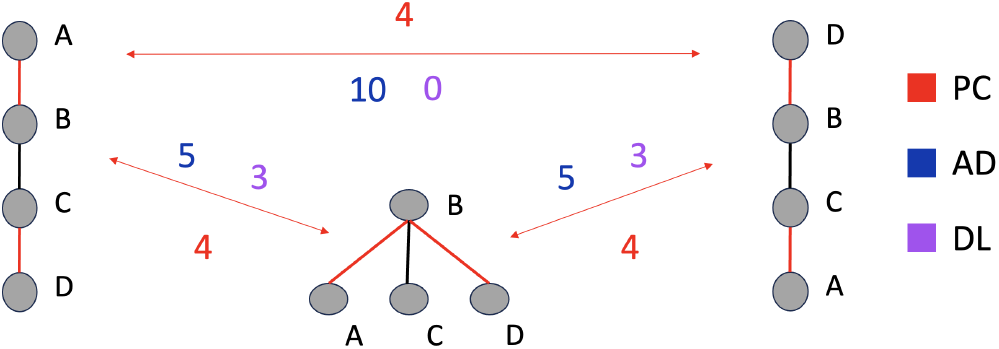
Three simple clonal trees (each labeled with mutations A-D) and their pairwise PC distances (each pair has a distance of 4 - indicated in red). Red edges indicate those edges that contribute to the PC distance. Pairwise AD (blue) and DL (purple) distances are included as well.

#### 2.2.2 Ancestor-Descendant distance

Ancestor-Descendant (AD) distance counts the number of pairs of ancestor-descendant mutations that exist in one tree but not in the other [12]. More formally, if *AD* (*T)* represents the set of ordered ancestor-descendant pairs observed in tree *T*, then the AD distance between two trees *T*_1_ and *T*_2_ is *AD* (*T*_1_, *T*_2_) =|*AD* (*T*_1_)△*AD* (*T*_2_)|, or the size of the symmetric difference (△) between the associated sets. In Figure 2 the left tree exhibits the following set of ordered ancestor-descendant pairs: {(*A,B*),(*A,C*),(*A,D*), (*B,C*), (*B,D*), (*C,D*)} and the middle tree exhibits the follow set of ordered ancestor-descendant pairs: {(*B, A*), (*B, C*),(*B, D*)}. The symmetric difference between these sets is {(*A, B*), (*A, C*), (*A, D*), (*C, D*), (*B, A*)}, so the AD distance between these trees is 5.

When multiple mutations appear on a single node, each mutation contributes individually to the associated ancestor-descendant relationships. For example, if mutations *B* and *C* appear together on a node that is a descendant of mutation *A*, they would contribute the following ancestor-descendant relationships: (*A, B*), (*A, C*). Furthermore, since clustered mutations indicate that the tree inference algorithm was unable to order those mutations, we include all pairs of clustered mutations as ancestral to each other. For example, if mutations *A* and *B* appear together on a node, we include the following ancestor-descendant pairs: (*A, B*), (*B, A*).

#### 2.2.3 Distinct-Lineage distance

Distinct-Lineage (DL) distance (or “Incomparable-Pair” distance) counts the pairs of mutations that are on distinct lineages in one tree (so neither mutation is ancestral to the other), but not in the other tree [7]. More formally, if *DL* (*T*)represents the set of unordered pairs of mutations that are on distinct lineages in tree *T*, then the DL distance between two trees *T*_1_ and *T*_2_ is *DL* (*T*_1_, *T*_2_) = |*DL* (*T*_1_)△*DL* (*T*_2_)|, or the size of the symmetric difference (△) between the associated sets. In Figure 2 the left tree exhibits no pairs of mutations that appear on distinct lineages (since the tree only contains a single lineage), but the middle tree exhibits the following set of distinct lineage pairs: {(*A, C*), (*A, D*), (*C, D*)}, so the DL distance between these trees is 3.

It is important to note that the DL distance is closely related to the AD distance. Specifically, a pair of mutations contribute to the DL distance when they are on distinct lineages in one tree, and are ancestral to each other in the other tree. This implies that the same pair of mutations will also contribute to the AD distance. A pair of mutations that are ancestral to each other one tree, and are ancestral to each other on the other tree, but in the opposite order, will contribute to the AD distance, but not the DL distance. As such, *DL* distance for a pair of trees will always be strictly less than the *AD* distance for that same pair of trees.

When multiple mutations appear on a single node, each mutation contributes individually to the associated distinct lineage relationships. For example, if mutations *B* and *C* appear together on a node that is a distinct lineage from mutation *A*, they would contribute the following distinct-lineage pairs: (*A, B*), (*A, C*).

## 3 Analysis of clonal tree distances

Most clonal tree distances are constructed to measure one or a few types of specific differences between a pair of clonal trees. For example, parent-child distance only measures direct connections between mutations. However, there are broader classes of important differences between clonal trees, of which it would be interesting to know how well existing distance measures capture or reflect these. In Section 3.1 we describe how we identified four potentially important types of broader changes that often appear in clonal trees: (1) lineage rearrangement, (2) subtree location, (3) cluster expansion, and (4) distinct mutations. Then in Section 3.2 we describe and analyze these four categories of tree difference, including how they impact the three distance measures described in Section 2.2.

### 3.1 Identifying broader tree difference categories

We identified broad categories of important types of changes that may appear when comparing pairs of clonal trees through conversations with several researchers who work directly in the field of clonal tree inference. In particular, we recruited a researcher to conduct a recorded exploratory session with an early prototype of our visualization tool (described in Section 4) so as to observe what features of the clonal trees they focused on (and get some initial usability feedback). Specifically, we presented the researcher with two clonal trees for a triple-negative breast cancer patient [26]. The clonal trees we used are those reported in [14] that were inferred by applying two clonal tree inference methods, PhiSCS [17] and SiFit [27], to single-cell sequencing data. The exact clonal trees used are those appearing in Figure 4 later in this paper. We prompted them to examine the trees to form insights about their comparison while providing a running commentary in an open-ended exploration task. We did not answer questions until a debriefing session at the end. Guided by the events that our interviewee identified in the session, and other discussions with experts in the field, we extracted a generalized set of important ways in which clonal trees might differ that go beyond the singular features captured by many distance measures.

### 3.2 Classes of tree difference

From our domain expert interviews, we identified four primary *tree difference classes*: common ways that two clonal trees might differ from one another that are particularly relevant on a broader scale. In this section, for each class, we discuss: (1) the definition of the class, along with an example of how it looks on a traditional node-link diagram; (2) possible biological implications of the difference class; and (3) how different existing tree distance measures might capture information about the difference class. We emphasize the importance of item (3) as knowing how a distance measure emphasizes these changes is essential to understanding when to use it over another option.

#### 3.2.1 Lineage rearrangement

A lineage rearrangement occurs when the order of mutation accumulation along a particular path from the root (i.e., along a lineage) changes from one tree to another. In Figure 3(a) we see a lineage rearrangement as mutation *G* is an ancestor of *D* in the tree on the left, but this relationship is reversed in the tree on the right. While in this example, the pair in question are directly adjacent as parent and child, lineage rearrangement can apply to swaps of mutations much further apart from one another, or may include nodes that contain multiple mutations.

**Figure 3:**
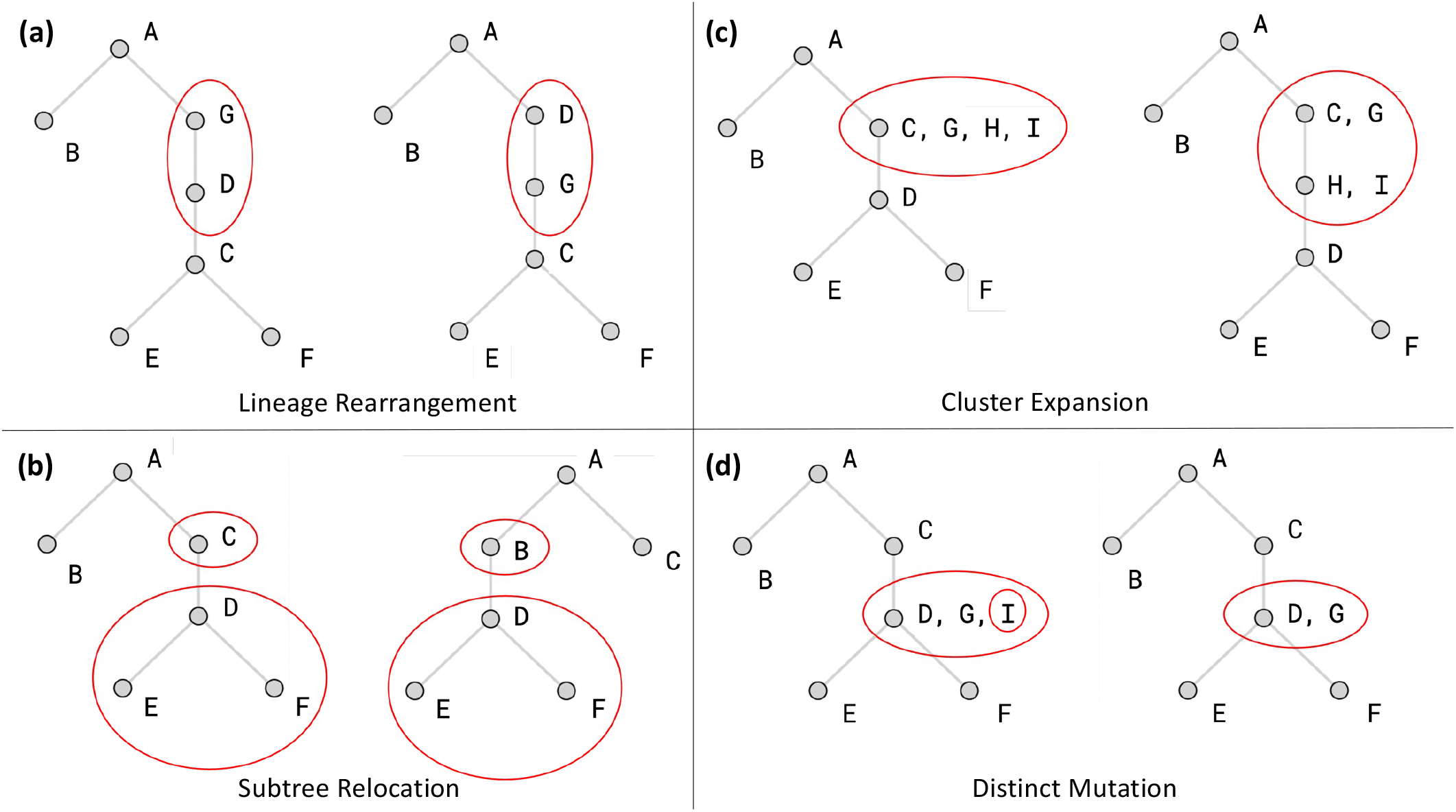
The four tree difference classes that we identified applied to example tree pairs. Nodes indicated with red circles are involved with the specified difference class.

**Figure 4:**
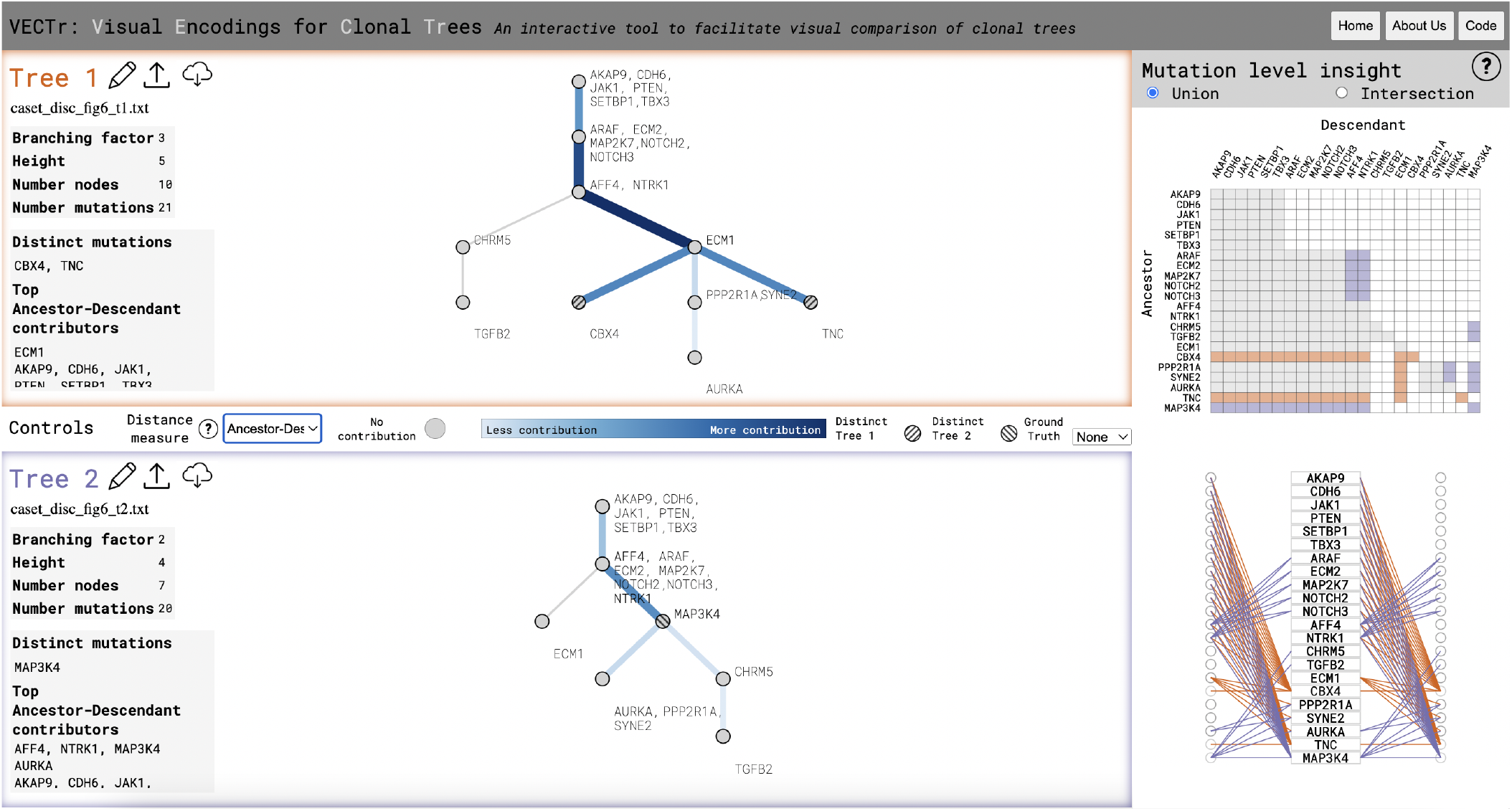
VECTr captures where differences occur between two trees inferred from a triple-negative breast cancer dataset [26].

Identifying instances of lineage rearrangement is important since the order in which mutations appear in a single lineage could impact their likelihood of being a “driver mutation,” – mutations that drive the growth of a tumor. Furthermore, the order of mutations may influence how many cells inherit which mutations, which can have an impact on how the patient might respond to treatments.

Distance measures react differently to instances of lineage rearrangement, though none seem to capture them directly. PC distance may be significantly impacted, depending on the size of the tree. The two mutations in question will certainly have different parents and children across the two trees. If there are at least three edges between them, a swap of this kind could add 8 or more to the PC distance scores (depending on the degree of the affected nodes). However, beyond this threshold of three edges, lineage rearrangements across greater distances will not have any additional impact on the distance measure. On the other hand, the impact of a lineage rearrangement on AD distance will depend on how far apart the rearranged mutations are from each other on the lineage. If the rearranged mutations are near to each other, the impact will be less, but if they are farther apart, all mutations that exist on the lineage between them will be impacted–having altered ancestor and descendant relationships with these mutations, thus increasing the measured distance. Figure 6 shows exactly that. In contrast, the DL distance will not capture this kind of change at all, since the mutations being swapped are on the same lineage and this distance only looks at mutations that are on different branches.

#### 3.2.2 Subtree relocation

A subtree relocation occurs when a subtree is re-located from one place in a clonal tree to another. Figure 3(b) shows an example of a subtree relocation where *DEF* moves from being a descendant of *C* in the tree on the left to being a descendant of *B* in the tree on the right. When this type of change happens, the ancestral relationships between all of the mutations in the moved subtree are preserved, but relationships between mutations in the moved subtree and the rest of the tree may change.

Biologically, a subtree relocation indicates that there is stronger evidence for how these mutations are related to *each other*, but less for how they relate to the *other* mutations. If any of the mutations in the relocated subtree are driver mutations, this could impact patients’ prognoses or even their treatment plan, depending on what (if any) druggable driver mutations exist in their tumor.

In terms of distance measures, PC distance will find little significance in this change. Given that the structure of the subtree stays the same (all the mutations are in the same place relative to each other), it will only find contribution to the distance at the nodes where the subtree is attached in both trees. In contrast, AD distance will weigh this kind of change more significantly as all of the mutations in the relocated subtree will acquire a different sets of ancestral relations and all previous ancestors will lose the mutations in this subtree as descendants. The DL distance will behave similarly to AD distance.

#### 3.2.3 Cluster expansion

A cluster expansion occurs when the evolutionary order of some set of mutations is ambiguous in one tree and ordered in the other tree–i.e., when a set of mutations are clustered into a single node in one tree, but separated out in the other. For example, in Figure 3(c) mutations *CGH I* belong to one node in the left tree, while on the right *CG* becomes ancestral to *H I* .

Specific ordering of a set of mutations has a potential impact on how those mutations are viewed as contributing to the tumor’s growth. For example, a mutation that occurred earlier in the tumor’s evolution–and thus closer to the root–may be more likely to be driving the growth of the tumor. When a set of mutations are clustered together, it can be more difficult to identify which mutation(s) are most likely to be contributing to the growth of the tumor.

When mutations are clustered on a single node, AD distance treats all clustered mutations as ancestral to each other. As such, when a cluster of mutations is split, we can now explicitly determine that some of these mutations are definitely ancestral to the others. So the magnitude of the effect of cluster expansion on AD distance depends on how many mutations become ordered. For PC distance, splitting the cluster causes some of the contributing mutations to obtain new parents and children, which can contribute to the total distance measured. DL distance will not capture this difference at all, as all of the mutations involved maintain their same lineage.

#### 3.2.4 Distinct mutations

A distinct mutation is a mutation that is present in one tree but not the other. For example, in Figure 3(d) mutation *I* is present in the left tree but not the right tree. This means that every relationship (e.g., ancestral, parental) involving this mutation is unique to the tree that contains the mutation.

Biologically speaking, knowing that there is an additional mutation that isn’t present in both trees could have a variety of implications, depending on the identity of the mutation and its location in the tree containing it. For example, the gene VHL is often mutated close to the root for many patients with kidney cancer [11]. If this mutation was missing from one of a pair of trees for a patient with this type of cancer, its absence from one tree is very notable.

As far as distance measures go, AD distance will be far more impacted by a distinct mutation than PC distance, especially as the size of the tree (in terms of number of mutations) increases. PC distance will only be locally concerned with the addition of a distinct mutation, with solely the mutations that are directly parents or children of the distinct mutation being affected. In contrast, in AD distance, every single mutation on the lineage of the distinct mutation will be impacted. DL distance will also be influenced broadly, with every mutation *not* on the lineage of the distinct mutation being impacted. Thus, the location of the distinct mutation as well as the number of branches in the tree dictates how AD distance and DL distance are affected.

## 4 Visualization of clonal tree distances

In the previous section we introduced 4 distinct classes of tree difference and analyzed how well these broad categories might impact a variety of distance measures. However, we relied on fairly simple node-link diagrams illustrate these features on only the simplest of data. While this analysis is useful in some cases, in real data the differences between two trees and how they contribute to a particular distance measure is not always so cut and dry. This seems like a clear candidate where specialized visualizations can be extremely useful. In the remainder of this section we introduce a tool called VECTr for visualizing clonal tree distances that utilizes three distinct visualization and describe how these visualizations are useful for highlighting instances of the four classes of tree difference described in Section 3.2.

### 4.1 VECTr: A tool for visualization of clonal tree distances

We have developed a web-based tool for visually comparing two clonal trees called *VECTr* (**V**isual **E**ncodings for **C**lonal **Tr**ees). VECTr allows users to upload a pair of clonal trees and visually explore their differences using three separate visual representations: (1) node-link diagram; (2) heatmap; and (3) tripartite, each described below. Users select a particular distance measure from Parent-Child, Ancestor-Descendant or Distinct-Lineage distance to guide and inform the notions of distance within their comparison task. Each visualization changes to reflect the choice of distance measure. Figure 4 shows an overview of what VECTr would display to a user when analyzing two clonal trees. VECTr was written using the D3.js library [3] and Flask.

### 4.2 Node-link diagram

VECTr’s central view is a pair of node-link diagrams representing the two compared trees. Figure 5 shows the encodings of these diagrams on simple examples, while Figure 4 shows it in context of the full tool. One of the motivations for creating VECTr was the fact that a standard node-link diagram has trouble highlighting all of the classes of tree difference articulated in Section 3.2. Nevertheless, it is the most familiar means of visualizing a clonal tree and provides a valuable reference for understanding patterns identified in VECTr’s other visualizations. We therefore chose to augment the nodes and links with addtional encodings that help to emphasize the features of our identified tree difference classes.

**Figure 5:**
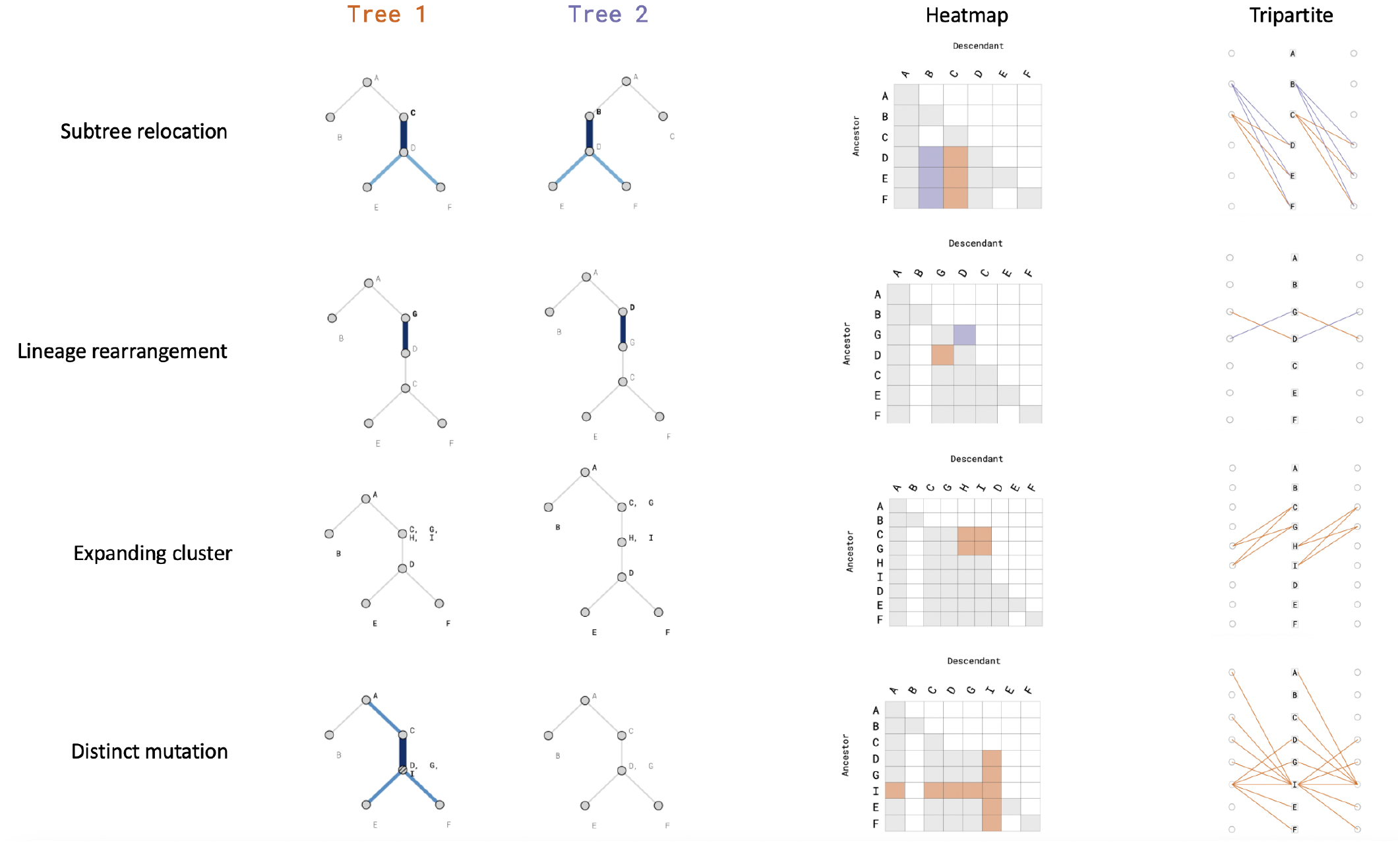
Examples of each of the four tree difference classes as shown by VECTr’s three visualization components.

First, we identify the mutation pairs that contribute to the differences between the two trees (as calculated by the selected distance measure) and highlight the edges between those mutations relative to that contribution. The contribution calculation for each edge is directly tied to the unique, relevant ancestral relationships involving that edge depending on the selected measure:

•**Parent-Child**. Edge weight directly corresponds to the number of PC pairs between the edge’s two incident nodes contributing to the total PC distance. For example, consider the edge *A, B → D*. If the PC pairs (*A, D*) and (*B, D*) contribute to the total PC distance, this edge will have a weight of 2.

•**Ancestor-Descendant**. Edge weight directly corresponds to the number of times that edge appears on *the path* between each mutation in all AD pairs contributing to the total distance. For example, consider the following arrangement of nodes: *A, B → C → D*. Suppose that only (*A, C*) and (*A, D*) contribute to the total AD distance. Then the edge *A, B → C* is weighted by 2.

•**Distinct-Lineage**. As discussed in Section 2.2.3, distinct lineage mutation pairs only contribute to the distance when the same mutations are ancestral to each other in the other tree. Therefore, we encode edge weights similarly to AD distance. Edge weight directly corresponds to how many times that edge appears on the path between each mutation in a DL pair, when looking at those mutations in the tree in which they are ancestral to each other.

We use width and color to highlight these edges, making it easier to identify the location and relevant mutations of structural differences between the two trees. This measure of contribution is also shown through the font-weight of the labels for each individual mutation, with mutations that contribute highly to the distance given a bolder font-weight. Finally, nodes containing distinct mutations (see Section 3.2.4) are highlighted with a hashed texture.

### 4.3 Heatmap visualization

To show how individual mutations contribute to distance, we employ a matrix heatmap visualization. Each possible pairwise relationship between mutations is signified by a square in the matrix, with rows representing ancestors and columns representing descendants. A filled box indicates that the specific relationship exists within solely Tree 1 (orange), Tree 2 (purple), or both trees (gray). The heatmap explicitly encodes every specific changed relationship between the two trees, affording an ease of lookup that is absent from the node-link visualization and allowing certain patterns of change to be readily identifiable through shape detection. Particularly, the heatmap provides further detail when interpreting differences between two trees as the result of a *cluster expansion*. As mutations within a cluster are all ancestral to each other when considering AD distance and DL distance, pulling a subset of mutations out of that cluster enforces a stricter ancestral order, leading to some mutations losing ancestors and others losing descendants. The heatmap captures these changing relationships better than the node-link diagram. Also, the heatmap handles the movement of groups of mutations well, as a purple and orange column of the same height is an easily recognized pattern (see *Subtree relocation* in 5). Another common shape is caused by a distinct mutation, which looks akin to a sideways T or L, given that all mutations on its lineage will gain a relationship.

### 4.4 Tripartite visualization

The third visual component of VECTr, which we call the *tripartite graph*, provides an alternative view of the ancestral/parental relationships between mutations (depending on the distance measure). To read it, users should look down the middle of the tripartite graph where there are mutation labels contained in rectangular nodes. Incoming edges on the left imply that the mutation has a unique ancestor/parent and outgoing edges on the right imply that the mutation has a unique descendant/child. Especially when there are few differences between the two trees, the tripartite graph reduces visual clutter and makes certain differences, like *lineage rearrangement*, more salient. In general, lineage rearrangement creates symmetrical patterns (Figure 6) while subtree relocation creates asymmetrical patterns. This effect becomes even more pronounced when the trees get larger and the distance of rearrangement becomes greater (see Section 5.1).

**Figure 6:**
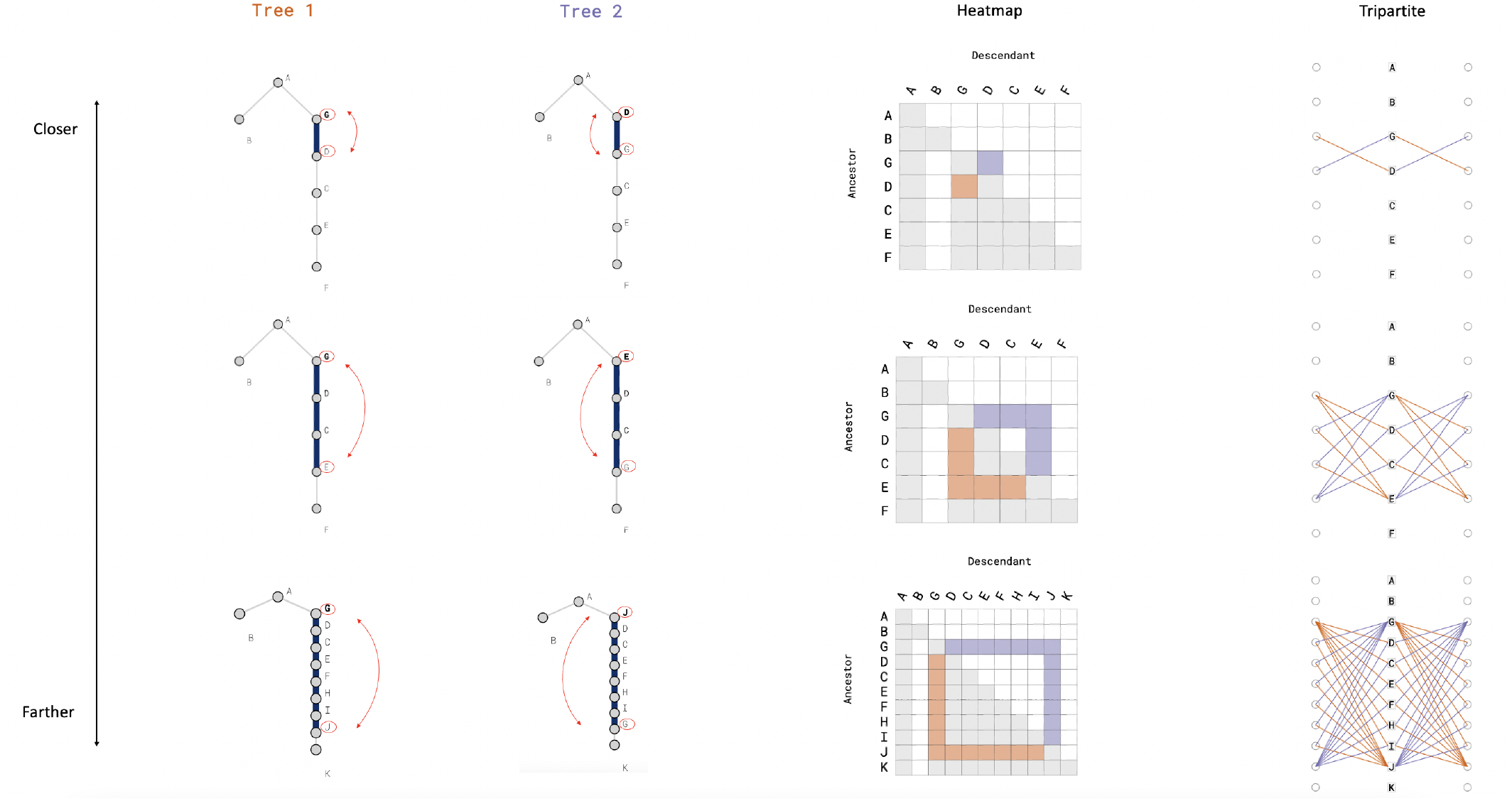
Iteratively increasing the distance between two swapped mutations on the same lineage.

### 4.5 Other features

In addition to the visualizations described above, we support several other features to aid interpretation.

### Summary statistics

To aid interpretation of tree structure, we provide summary statistics in a column to the right of the node-link diagrams (see Figure 4), including branching factor, height, and node and mutation counts. Additionally listed are the distinct mutations for each tree (see Section 3.2.4) as well as those that are the “top contributors” to the selected distance measure.

### Interactivity

Brushing over a mutation in any of the views within VECTr initiates linked highlighting of that mutation across all views (or that *set* of mutations when hovering over a node). This helps connect insights relevant to tree difference classes that may be easier to identify in one visualization versus another. Clicking on the name of a mutation will open that mutation’s page in the GeneCard’s gene database [24], making it easier for domain information to inform insight and analysis in VECTr.

### Union/intersection of mutations

Since distinct mutations in one tree can end up dominating the contributions to distance and visually drowning out more nuanced changes, users can decide to only show *non-*distinct mutations (i.e., the intersection of mutations across the two trees) in the visualizations on the right.

### Ground truth annotation

While often researchers may want to compare two plausible trees inferred using different algorithms from the same data, sometimes there may be a definitive “ground truth” (e.g., when inference methods are being tested on simulated trees). In such instances, users can assign one tree to be ground truth. This preserves all of the ancestral relationships between mutations in that tree to be “correct,” which then highlights all the manners in which the other tree strictly differs. Labeling one tree as ground truth shifts VECTr’s color scheme from purple/orange for the two trees to simply encoding any departures from the ground truth in the other tree using a red map (see Figure 7).

**Figure 7:**
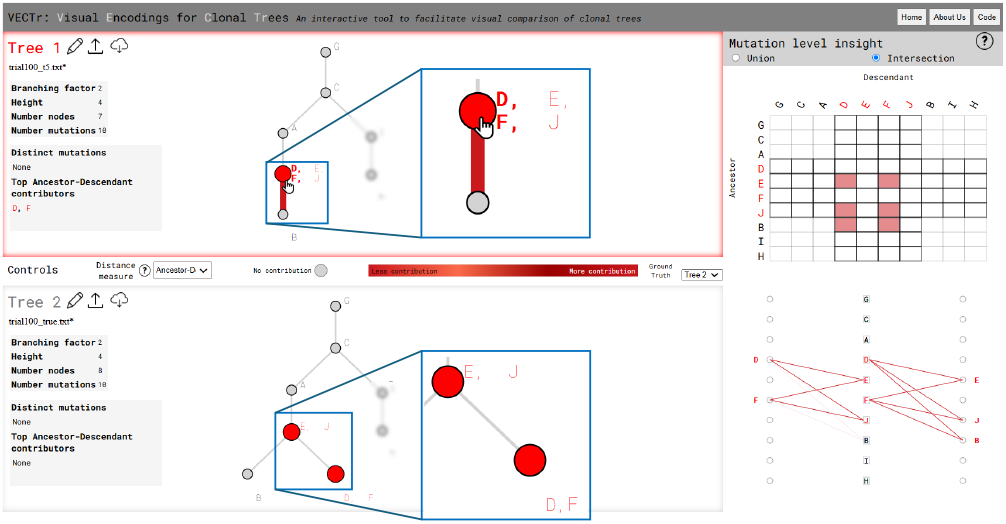
A full view of the VECTr interface when comparing two clonal trees when Tree 2 is designated as the “groundtruth” tree. Zoom-in annotations are provided for the nodelink diagram.

### Upload/download/modification of trees

Trees can be uploaded to VECTr using the common Newick file format and/or input or modified directly in VECTr. The ability to dynamically edit trees within a session is helpful for inputting custom trees from scratch, modifying naming conventions, or observing how small changes might alter differences across the two trees. Trees can be downloaded as SVG, making for easy figure generation.

## 5 Use cases

To illustrate the utility of our tool, we lay out three use cases. The first use case (Section 5.1) discusses a general example of how our heatmap and tripartite visualizations can help users identify a specific class of tree difference more easily than is possible with a standard node-link. The second use case shows a more specific instance of using our tool when a “ground truth” tree is available. Finally, the third case walks through a real-data example of comparing two trees exhibiting multiple classes of tree difference, showcasing the interactive workflow afforded by our tool.

### 5.1 Identifying tree difference classes

Consider the three cases of *lineage rearrangement* in Figure 6, in which the distance between the two swapped mutations increases from 1 edge, to 3, to 7 from top to bottom. While the annotated node-link diagrams highlight the entire lineage, this can be deceptive, as apart from the swapped mutations, the *intermediate* lineage in the chains actually remains unchanged. However, the heatmap provides a distinct pattern with this event. The “cornerless square” shape clearly identifies the two mutations swapped and which direction they moved in the tree. Similarly, we see a distinct “butterfly” shape in the tripartite graph (or at least that’s how we came to describe it). The butterfly shape affords the same clear narrative about which two mutations were involved, with densely woven “wings” originating from these mutations.

For complex trees exhibiting multiple, potentially overlapping classes of tree difference, these patterns retain visual saliency (see an example in Section 5.3). Users can come to quickly identify these patterns through perceptual mechanisms of shape recognition, more quickly attuning to them than would be possible by scouring the node-link view mutation by mutation.

### 5.2 Ground-truth comparison

Next, we consider comparing a “ground-truth” tree to one that has been produced using a new inference method (such as is commonly done when benchmarking new clonal tree inference methods). Consider a scenario where a researcher wants to evaluate the performance of their novel clonal tree inference method. They first create (or obtain) a simulated dataset with known ground-truth clonal tree reconstructions. They also apply their method to the simulated data, yielding its own set of tree reconstructions. Suppose the researcher uses Ancestor-Descendant (AD) distance to compare their inferred clonal trees to the ground-truth trees. They might see that the distance between the inferred trees and ground-truth trees is generally small, which is promising, but they need a more nuanced understanding of what the differences are. They would then load a tree from their novel method and the ground truth tree into VECTr (see Figure 7). The novel tree’s departures from the ground-truth are annotated in red, and the first thing that might draw attention is a single highlighted edge. It is connected to a node with two mutations (*D* and *F*) in a heavier font weight, indicating they are top-contributing mutations. Hovering over this node activates linked highlighting across the node-link diagram, heatmap, and tripartite graph. Looking down the middle of the crisscrossed edges in the tripartite graph provides an explanation for why mutations *D* and *F* are top contributors: they are involved in 6 unique AD pairs. The researcher might now form these hypotheses: (1) those mutations have moved together up their lineage in Tree 1 when compared to Tree 2; or (2) the mutations have relocated to another subtree entirely. Referring back to the node-link, our researcher concludes that hypothesis (1) was correct as *D* and *F* are both clustered as a leaf node in Tree 2 but move up their lineage to join mutations *E* and *J* to form a larger cluster. This process can then be repeated for additional trees to see if patterns across the reconstructions emerge.

### 5.3 Real data comparison

For our final use case, we examine two clonal trees inferred from data for a colorectal cancer patient [16]. In this instance, the two trees were inferred using the same inference method, but while allowing different assumptions about the underlying data [17].

Using VECTr, a researcher might quickly notice “driver mutations” which have known biological importance in cancer. For example, *TP53* (the most commonly mutated gene in human cancers [4]) is near the top of both trees in Figure 8. They can easily obtain additional information about this or other potential driver genes by clicking on their labels to see their GeneCards.

**Figure 8:**
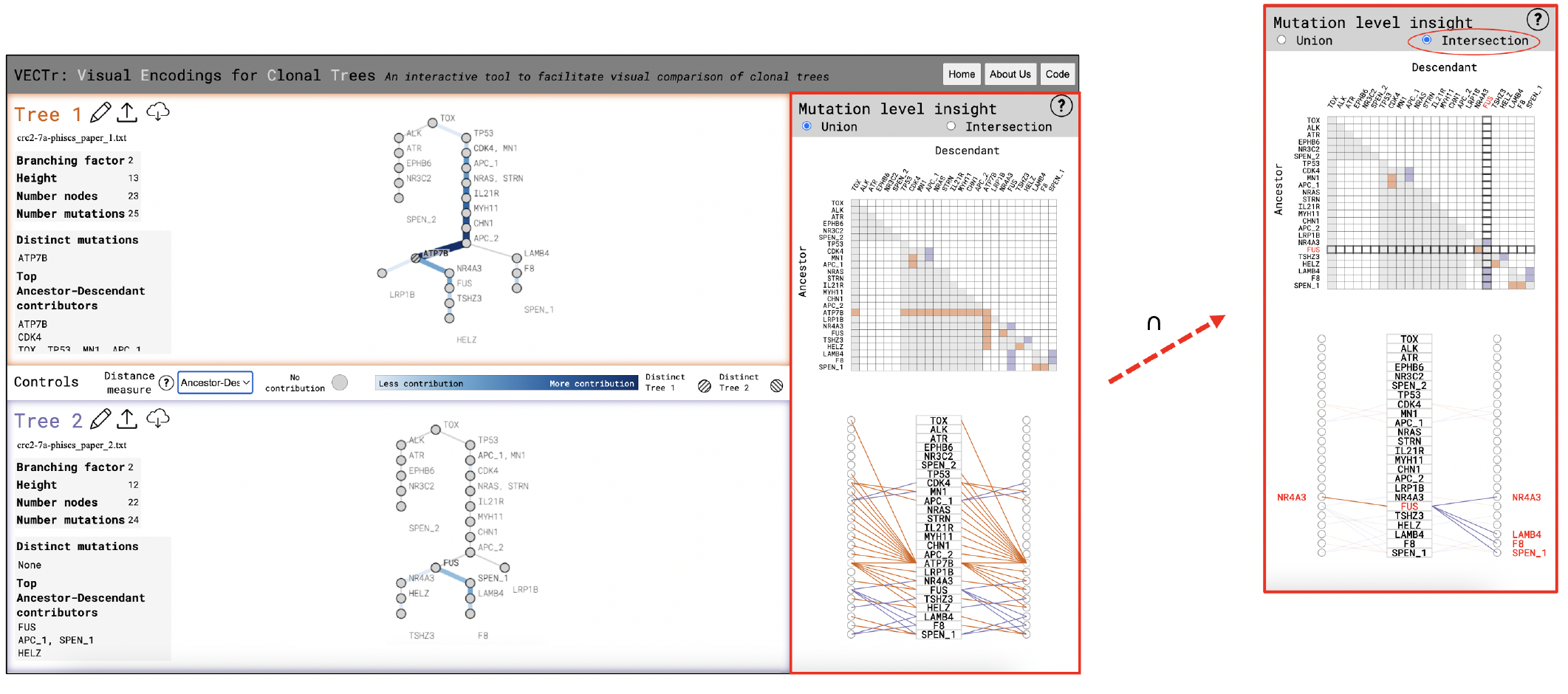
A full view of the VECTr interface when analyzing real data. Both clonal trees are inferred from a colorectal cancer patient from [16]. PhISCS [17] inferred both clonal trees but with different parameters. The callout depicts the updated heatmap and tripartite after selecting the “intersection” option. There, FUS is highlighted via mouseover, along with each mutation with which it has a changing relationship: NR4A3, LAMB4, F8, and SPEN_1.

Using Ancestor-Descendant (AD) distance, the distinct mutation *ATP7B* jumps out. However, it dominates the visualizations, particularly the heatmap and tripartite (see Figure 8). After noting its presence, the researcher selects the option to show only the *intersection* of mutations in the visualizations on the right (described in Section 4.5). This removes the distinct mutation from the mutation-level visualizations so that the relationships between other shared mutations become more clear (see callout in Figure 8).

The tripartite visualization shows two local sets of differences. The top set (with *CDK4, MN1*, and *APC_1*) seems reminiscent of the “butterfly” shape discussed in Section 5.1 (and seen in Figure 6), suggesting a possible *lineage rearrangement* event. Honing in on the three mutations, it becomes clear that *CDK4* and *APC_1* swapped ancestral order. Additionally, a *cluster expansion* event occurred related to the cluster with MN1. The movement of *CDK4* and *APC_1* is high on the tree, indicating that these mutations occurred relatively early in the development of the tumor.

The bottom set of differences (involving *NR4A3, FUS*, etc.) contains more variation across relationships and thus is more complex to parse. However, by examining a few specific mutations, the individual movements can be untangled and identified. For example, the researcher might first inspect *FUS* by mousing over its label, as shown in Figure 8. This highlights its location in both trees and its context among other mutations in the heatmap and tripartite visualizations. The specific pattern around *FUS* in the tripartite is mostly asymmetric (excepting *NR4A3*), with *LAMB4, F8*, and *SPEN_1* all only connected on the right side in purple (meaning Tree 2). Now, the researcher can determine that *FUS* rearranged its order with *NR4A3*, as well as gained the *LAMB4, F8*, and *SPEN_1* subtree (that had branched from *APC_2* in Tree 1). A better understanding of the differences here may help the researcher to assess if one reconstruction provides a simpler or more logical explanation for how that process may have occurred.

## 6 Conclusion

In this paper, we provide important analysis of several existing distance measures for clonal trees. Specifically, we define a set of four classes of tree differences that describe common ways that clonal trees might differ from each other and analyze these classes using three different clonal tree distance measures. We also present a novel visualization tool, *VECTr*, for use in conjunction with these distance measures. We find that our tool which incorporates several novel visualizations for interpreting clonal trees - provides more nuanced information than just the application of a distance measure alone. Furthermore, our tool allows for enhanced identification of our identified tree classes. Finally, we demonstrate the utility of our work using both simulated and real data.

Key areas for future work include the incorporation of a broader set of distance measures and expanding the potential for comparisons beyond single pairs of clonal trees, potentially out into “neighborhoods” of similar trees inferred from real data or synthetically generated.

## Acknowledgments

This work has been supported by the National Science Foundation (NSF) award CAREER-IIS-2046011.

